# FORMATION OF LIPOFUSCIN-LIKE AUTOFLUORESCENT GRANULES IN THE RETINAL PIGMENT EPITHELIUM REQUIRES LYSOSOME DYSFUNCTION

**DOI:** 10.1101/2021.02.23.432539

**Authors:** Cristina Escrevente, Ana S. Falcão, Michael J. Hall, Mafalda Lopes-da-Silva, Pedro Antas, Miguel M. Mesquita, Inês S. Ferreira, M. Helena Cardoso, Ana C. Fradinho, Clare E. Futter, Sandra Tenreiro, Miguel C. Seabra

## Abstract

**Purpose:** We aim to characterize the pathways required for autofluorescent granule (AFG) formation by retinal pigment epithelium (RPE) cells using cultured monolayers.

**Methods:** We fed RPE monolayers in culture with a single pulse of photoreceptor outer segments (POS). After 24h the cells started accumulating AFGs similar to lipofuscin *in vivo*. Using this model, we used a variety of light and electron microscopical techniques, flow cytometry and western blot to analyze the formation of AFGs. We also generated a mutant RPE line lacking Cathepsin D by gene editing.

**Results:** AFGs appear to derive from incompletely digested POS-containing phagosomes and are surrounded after 72h by a single membrane containing lysosome markers. We show by various methods that lysosome-phagosome fusion is required for AFG formation but that impairment of lysosomal pH or catalytic activity, particularly Cathepsin D activity, enhances AF accumulation.

**Conclusions:** We conclude that lysosomal dysfunction results in incomplete POS degradation and AFG accumulation.

## Introduction

Age-related macular degeneration (AMD) is the most common blinding disease in the western world, characterized by loss of vision in the macular (central) area of the retina, which impacts the quality of life of the elderly. Currently, there are no effective therapies for the common forms of AMD namely early, intermediate or late stage “dry” AMD, also called geographic atrophy ^1, 2^. The primary cause of pathology in AMD appears to be retinal pigment epithelium (RPE) degeneration. RPE thinning and depigmentation leads to atrophy and the accumulation of extracellular deposits called drusen between the RPE and the choroid ^3^. Eventually, RPE atrophy leads to photoreceptor loss. In fact, photoreceptors, responsible for phototransduction, rely on the RPE for nutrients and waste disposal.

The RPE is responsible for the daily digestion of photoreceptor outer segments (POS), a process essential for sustained photoreceptor function which requires the regular recycling of visual cycle components ^4^. However, POS digestion by RPE cells imposes a continuous heavy burden on the lysosomal network of these non-dividing cells. Another AMD hallmark is the accumulation of lipofuscin in the RPE ^5, 6^. The appearance of lipofuscin, clinically detected as fundus autofluorescence occurs naturally and accumulates gradually with age but it is exacerbated in disease, being a predictive marker of retinal stress ^7–9^.

The age-related pigment lipofuscin is particularly abundant in nerve cells, cardiac muscle cells and skin. Accordingly, its origin as well as its composition vary among tissues, being mainly constituted by oxidized proteins (30–70%), lipids (20–50%), metal cations (2%), and sugar residues ^10^. These differences in composition are reflected in very wide spectra of lipofuscin fluorescence emission ranging from 400 to 700 nm ^11^. Lipofuscin accumulates in lysosomes and for many cell types, is thought to originate internally resulting from altered cellular proteostasis, autophagy and mitophagy and/or dysfunction of lipid metabolism ^10^. Lipofuscin in the RPE is proposed to be a by-product of POS digestion and the final maturation step of ingested phagosomes is fusion with lysosomes ^8, 12–14^.

Lysosomes are critical players in the complex cellular network that comprises endo-lysosomal, phagocytic and autophagic pathways, coupled with metabolic sensing ^15^. Lysosome involvement in phagosome degradation within the RPE was initially described by EM studies that identified acid phosphatase activity in both lysosomes and POS phagosomes ^13, 14^. Over the years, multiple studies have demonstrated that POS phagosome degradation is a very complex multistep process, involving different hydrolases and the coordination of different molecular motors (reviewed in ^4^).

The heavy, daily, phagocytic load on the RPE, together with a highly oxidative environment and high metabolic activity, leads to the slow accumulation of indigestible material over decades. The inability to fully digest POS and the lifelong accumulation of undigested material led to the idea that lysosomal dysfunction in the RPE is at the centre of a pathogenic hub contributing to proteotoxicity, mitochondrial dysfunction, redox imbalance and inflammation in AMD, eventually leading to RPE cell death and vision loss ^16^. More generally, evidence indicates that the lysosomal network is prone to age and disease-related dysregulation, including in neurodegenerative diseases like Parkinsońs Disease ^17, 18^.

The formation and maintenance of lipofuscin within RPE cells remains ill-characterised. In this work we have optimized an *in vitro* model system that recapitulates AMD features. Feeding RPE cells with a single pulse of porcine POS leads to accumulation of autofluorescent granules (AFGs) similar to lipofuscin *in vivo* ^19–25^. Our results suggest that some lysosome activity, in particular lysosome-phagosome fusion is required for AFG formation but that impairment of lysosome catalytic activity, particularly Cathepsin D (CTSD) activity, enhances AFG accumulation. Therefore, dysfunctional lysosomal activity leads to reduced POS degradation and increased AF accumulation.

## Methods

### Cell Cultures

Primary human fetal RPE cells (hfRPE) were purchased from Lonza and cultured using optimized RtEBM^TM^ Basal Medium and RtEGM^TM^ SingleQuots^TM^ Supplements (Lonza), according to the supplier’s instructions. For confocal or electron microscopy assays, cells were seeded on laminin-coated (BioLamina) coverslips (10 µg/mL). Cells were used after 21 days in culture, after which they demonstrate a cobblestone morphology with the formation of tight junctions confirmed by transepithelial electric resistance (TEER) values of ± 250 Ω.cm^2^, measured using an electrode (STX2; World Precision Instruments). ARPE-19 cells (ATCC) were cultured in Dulbecco’s Modified Eagle Medium/Nutrient Mixture F-12 (DMEM/F-12) (Gibco) supplemented with 10% fetal bovine serum (FBS) (Gibco) and 1% Penicillin-Streptomycin (Pen-Strep) (Gibco). For porcine RPE (pRPE) primary cultures, porcine eyes were collected from a slaughterhouse, kept on ice and the procedure for cell culture was performed on the same day. Briefly, external tissue from the eyeballs was removed and eyes were cleaned in an iodine surgical scrub solution, diluted 1:4 in water. Eyes were washed in a PenStrep/PBS solution for 5 min. The eyes were opened, inside the tissue culture hood, using a scalpel, and kept in multiwell dishes, opening facing up. The eye cups were filled with PBS and the neural retinas were removed. PBS was replaced by trypsin and the eye cups were incubated for 30 min, at 37°C. RPE was resuspended in Dulbecco’s Modified Eagle Medium: Nutrient Mixture F-12 (DMEM/F-12; Gibco) supplemented with 1% sodium pyruvate, 1% non-essential aminoacids (both from Biowest), 1% PenStrep and 1% FBS, spun at 1000rpm at room temperature, 5 min. The pelleted cells were resuspended in fresh medium (containing 10% FBS) and seeded in 6-well plates. This medium is changed once a week until cells reach confluency, at which point they are changed to reduced-serum medium (1% FBS). Primary cultures can be expanded and used for assays, by plating onto multiwell plates. Whenever using glass coverslips or transwells, cells were seeded on laminin-coated (BioLamina) coverslips (10 µg/mL). All of the cell models were grown in a 5% CO_2_ incubator at 37 °C.

### POS Isolation and Gold Tagging

Photoreceptor outer segments (POS) were isolated from porcine eyes, according to ^26^ with some minor modifications. Briefly, porcine eyes were collected from a slaughterhouse and kept on ice. External tissue from the eyeballs was removed and eyes were washed in a 1% Penicilin/Streptomycin (PenStrep; Gibco) in phosphate buffer saline (PBS) solution for 5 min. The eyes were opened, using a scalpel, and kept in multiwell dishes, opening facing up. The eye cups were filled with PBS and the neural retinas were removed and collected in a falcon containing homogenization solution (20% sucrose, 20mM tris acetate pH 7.2, 2mM MgCl_2_, 10mM glucose, 5mM taurine). Retina homogenate was filtered through gauze and added to a continuous sucrose gradient (25-60% sucrose, 20mM tris acetate pH 7.2, 10mM glucose, 5mM taurine). POS were separated by centrifugation at 25.000 rpm (76.7g) for 120 min, in a swing rotor (SW-32-Ti; Beckman). A single orange band in the upper third of the gradient, corresponding to POS, was aspirated with a P1000 tip and collected into a new falcon. POS were washed three times (wash 1: 20mM tris acetate pH 7.2, 5mM taurine; wash 2: 10% sucrose, 20mM tris acetate pH 7.2, 5mM taurine; wash 3: 10% sucrose, 20mM sodium phosphate pH 7.2, 5mM taurine) by sequentially centrifugations (5000 rpm for 10 min at 4°C) and resuspension of the pellet. POS preparations were washed in PBS and resuspended in medium and protein content was measured (BCA Protein Assay Kit, Thermo Scientific) and in parallel, POS particles were counted in a cell counting chamber POS then were aliquoted and stored at −80° C, in medium containing 10% fetal bovine serum (FBS; Gibco), 2.5% sucrose, 0.04% sodium azide and 1% PenStrep. For the UV-irradiated POS, each prep was exposed to ultraviolet radiation using a UVP Crosslinker (CL-1000 Model, Analyticjena) with 2×2 min pulses of 254 nm at an estimated radiant exposure of 1 J/cm^2^. For gold-tagging, POS preps were thawed, pelleted and re-suspended in 200μl 10 nm gold colloid solution (BBI Solutions). After a mild sonication for 10 min in a sonicator bath, POS were washed three times in PBS.

### POS Phagocytosis Assays

Before the phagocytosis assays, POS preps were thawed in the dark, washed three times in PBS by spinning for 10 min at 5000 rpm. For the UV-irradiated POS, each prep was exposed to ultraviolet radiation using a UVP Crosslinker (CL-1000 Model, Analyticjena) with 2×2minutes pulses of 254 nm at an estimated radiant exposure of 1 J/cm2. RPE cells were fed during 4h with POS at a final concentration of 50 (6.5×10^4^ POS particles/cm^2^) or 200 µg/mL (∼2.6×10^5^ POS particles/cm2) in Dulbecco’s Modified Eagle Medium: Nutrient Mixture F-12 (DMEM/F-12; Gibco) supplemented with 1% sodium pyruvate, 1% non-essential aminoacids (both from Biowest), 1% PenStrep and 10% FBS. After this pulse (time 0h), POS were removed, cells were washed and cultured in supplemented DMEM with 1% FBS during 4h, 24h, 3 and 7 days. After these chase periods, cells were either fixed for confocal/electron microscopy studies or trypsinized for flow cytometry assays.

### Cell Treatments

After the 4h POS feeding, cells were treated during the 3 days chase period with the following compounds: Bafilomycin A1 (5-10 nM; Sigma-Aldrich), Rab7 inhibitor (CID 1067700; 5-25 µM; Sigma-Aldrich), Leupeptin (10 or 25 µM; Calbiochem) plus Pepstatin A (25 µM; Sigma-Aldrich) or a BACE 1 inhibitor (PF9283; 3-9 µM; Sigma-Aldrich). After the chase period, cells were trypsinized for flow cytometry assays.

### CRISPR/Cas9 Genome Engineering

To generate *CTSD* knock-out ARPE-19 cells, sgRNAs were designed for specific target sequences, as previously described ^27^. During sgRNA cloning the pSpCas9(BB)-2A-GFP (pX458) was used (a gift from Feng Zhang, Addgene plasmid #48138, http://n2t.net/addgene:48138, RRID: Addgene_48138). *CTSD* was targeted using the gRNA: ACGTTGTTGACGGAGATGCG. ARPE-19 cells were then transfected using Lipofectamine^TM^ 3000, according to the manufacturer’s instructions. Single colonies were grown and expanded for genomic DNA extraction. Indel mutations were confirmed by sequencing, using the primers: forward- 5 ’-TTTCTCTGTGCTGCCGCTTA -3’, reverse- 5’ - CATCGCAGCCAAGTTCGATG -3’. Genotyping revealed homozygous frameshift insertion (g.616_617insA).

### Flow Cytometry

For flow cytometry assays, cells were trypsinized with TrypLE™ Express Enzyme (Gibco) for 30 min. Cells were resuspended in DMEM supplemented with 10% FBS, washed two times with PBS and one time with flow cytometry buffer (1% FBS and 2 mM EDTA in PBS). Finally, cells were resuspended in flow cytometry buffer and data acquisition was performed in a FACS CANTO II flow cytometer (BDBiosiences) using the 488 nm excitation wavelength to evaluate cellular AF. At least 30,000 cells were acquired per condition using BD FACSDivaTM software (Version 6.1.3, BD Biosciences). Data analysis was performed in FlowJo (Version 10, BD Biosciences) and GraphPad Prism (Version 7). Results were represented as the percentage of 488-positive cells and were normalized to the values of AF detected in cells without POS pulse or cells pulsed with POS but in the absence of compounds.

### Confocal Immunofluorescence Microscopy

Cells grown on coverslips were fixed for 15 min in 4% paraformaldehyde (Alfa Aesar) or 100% methanol, at room temperature or −20°C, respectively. Cells were blocked/permeabilized for 30 min in PBS containing 1% BSA and 0.05% saponin or 0.1% TX100, according to the antibodies used. Cells were then incubated with primary antibodies, namely, mouse anti-RetP1 (ThermoFisher) (1:100), mouse anti-LAMP1 conjugated with Alexa fluor 647 (clone H4A3, BioLegend) (1:500), rabbit anti cathepsin D (Abcam), or Phalloidin 488 (ThermoFisher) for 1h, followed by incubation with Alexa-conjugated secondary antibodies donkey anti-mouse or anti-rabbit 647 (1:1000) (Invitrogen). Cell nuclei were labelled with DAPI (Sigma) (1 µg/ml) and cells were mounted in Mowiol mounting media (Calbiochem). AF was visualized in the 405 nm excitation wavelength. Images were acquired in a Zeiss LSM 710 confocal microscope or in a Zeiss LSM980 airyscan confocal in the Multiplex SR-4Y imaging mode, with a Plan-Apochromat 63×1.4 NA oil-immersion objective. Digital images were analysed using LSM Image software or ImageJ. AFGs and RetP1-positive POS were quantified using ImageJ software (https://imagej.nih.gov/ij/). Ten random images of each time point were acquired using a 63x magnification, with zoom 1. Alexa-fluor 488 Phalloidin was used to visualize cell limits and a Z-projection of all the slices containing cellular staining was done. The same threshold levels were applied to all images. AFGs were visualized in the 405 nm excitation wavelength and counted using ‘Cell counter’ plugin in ImageJ. Intracellular RetP1-positive POS labelled with an Alexa Fluor 647 were visualized and counted using the 633 nm excitation wavelength, to minimise interference from the AF signal since in this wavelength there is less AF signal. RetP1-positive aggregates with large dimensions and found attached to the outside of the cells were excluded from the counting. Results are represented as the average of the number of AFGs detected per field of view ± SEM. The percentage of intracellular AFGs surrounded by LAMP1 or CTSD was calculated as the number of AFGs surrounded by those markers/number of total intracellular AFGs observed per field of view ± SEM.

### Electron Microscopy

For correlative electron microscopy (CLEM) and transmission electron microscopy (TEM), cells were seeded on laminin-coated photoetched gridded coverslips (MatTek Corporation) and at the indicated times of POS chase, cells were fixed with 4% PFA (TAAB Laboratory Equipment Ltd) in PBS for 30 mins before image acquisition using an inverted Zeiss LSM710 (63x lens, NA 1.3, oil immersion). Samples were subsequently washed with PBS, fixed with 2% PFA, 2% Glutaraldehyde (TAAB Laboratory Equipment Ltd) in 0.1M Sodium Cacodylate for 30 minutes, osmicated and further processed for resin embedding^28^. Resin blocks were sectioned (Leica Microsystems UC7) and 70-nm ultrathin serial sections collected on formvar-coated slot grids were stained with uranyl acetate and lead citrate and observed with a transmission electron microscope, Tecnai G2 Spirit (FEI) or observed with a JEOL 1010 or a JEOL 1400 Plus transmission electron microscope and imaged with an Orius SC1000B charge-coupled device camera with Digital Microgaph software (both Gatan). EM and light microscopy data sets were registered manually in Fiji (ImageJ) and Power Point (Microsoft Office 365 ProPlus) using DIC images and serial EM images, where nuclear and plasma membrane features, together with lipid droplets were used as unbiased fiducials

### Western Blot

Cells were lysed in ice cold cell lysis buffer (Cell Signaling Technology) supplemented with protease and phosphatase inhibitor cocktails (Roche) according to the manufacturer’s instructions. Lysates were pelleted for 15 min at 13 000 xg at 4 °C and supernatants kept for protein quantification (BCA Protein Assay Kit, Thermo Scientific). Equal amounts of cellular proteins were resolved on 10% or 12% sodium dodecyl sulfate – polyacrylamide gels (SDS-PAGE) and subsequently transferred to nitrocellulose membranes (Bio-Rad Laboratories). Membranes were blocked using 5% non-fat dry milk or 5% bovine serum albumin (BSA) (Sigma-Aldrich) for phosphorylated proteins immunoblots in Tris-buffered saline (TBS) (50 mM Tris, 150 mM NaCl, pH = 7.6) containing 0.1% Tween-20 (Sigma-Aldrich) (TBS-T) for 1 h and primary antibodies were then added in blocking solution. The following antibodies were used: mouse anti rhodopsin 1D4 (Abcam) and mouse anti-β-Actin (Sigma-Aldrich). Primary antibody incubations were carried out at 4°C overnight. After washing with TBS-T, the appropriate HRP-conjugated secondary antibody was added (1:5000 in blocking buffer) for 2 h at room temperature. Antibody binding was detected using chemiluminescence ECL Prime Western Blotting Substrate (GE Healthcare).

### High Content Imaging

For high-content confocal imaging, cells were cultured on 96-well skirtless CellCarrier plates (PerkinElmer), and fixed using 4% PFA in PBS. Each experimental condition was repeated in 4 wells. Using an Opera Phenix high-content screening system (PerkinElmer), the same 3 randomly selected regions were imaged in each well, and 8 confocal slices separated by 1μm were acquired. For semi-automated analysis of the high volume of images acquired by high-content imaging, macros were designed in ImageJ to batch process all images identically. For quantitating the number of DAPI-stained or AF punctae, z-stacks were combined into a maximum intensity projection, before identifying the individual objects using the ‘Find Maxima’ tool. The appropriate tolerance value for the ‘Find Maxima’ tool was applied to all images for a single staining and was subjectively pre-determined using a sample of randomly selected images.

### Statistics

All our results are shown as mean ± SEM. Statistical significance within groups was assessed either by Student *t* test or one-way ANOVA followed by multiple comparisons Bonferroni or Dunnet post hoc correction, as appropriate, using GraphPad Prism, version 7 Software. p values less than 0.05 were considered statistically significant.

## Results

### One Single Feeding of POS Leads to the Formation of Autofluorescent Granules (AFGs)

To model the appearance of autofluorescence (AF) in the RPE and study its role in disease, we optimized an *in vitro* model whereby primary human fetal RPE (hfRPE) cells grown for 3 weeks were challenged with a single pulse of purified porcine POS (Fig. S1A) at two different concentrations, 50 (∼6.5×10^4^ POS particles/cm^2^) and 200 µg/mL (∼2.6×10^5^ POS particles/cm^2^). While most previous reports studying phagosome processing utilize POS labeled with a fluorescent probe and follow its fluorescence, we use unlabeled POS and follow the appearance of AF.

We used flow cytometry to quantify AF levels as described previously ^29, 30^. Immediately after POS feeding (0h), no AF is detected, comparable to the cells in the absence of POS (Fig. 1A). AF starts to be detected 24h after POS feeding and progressively increases with time, at least until 7 days later. Moreover, cells fed with lower POS concentrations (50 µg/mL) showed lower AF levels, suggesting that AF is dependent on the POS concentration used. Similar results were obtained when ARPE-19 monolayers (Fig. S1B) or primary pig RPE (pRPE) (Fig. S2A-C) were treated with a single pulse of POS and AF quantified by flow cytometry and high content screening, respectively.

**Figure 1.**
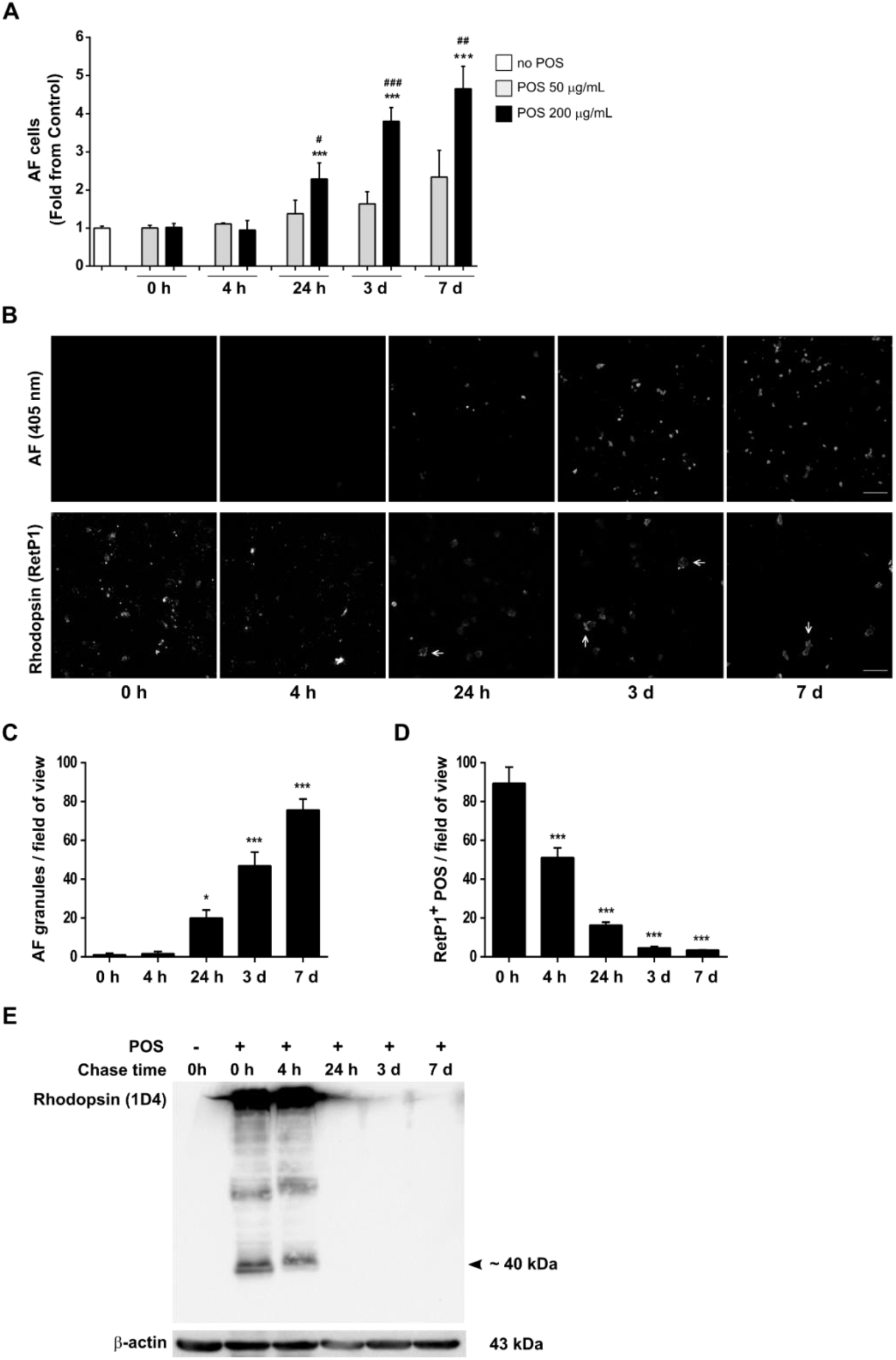
Lipofuscin-like AF formation in hfRPE cells. hfRPE cells pulsed with different concentrations of POS were monitored at the indicated time points after POS incubation. (**A**) AF levels were quantified flow cytometry as in “Material and Methods” and are represented as mean ± SEM of at least 3 independent experiments. Statistical comparison was performed using one-way ANOVA followed by Dunnet’s or Bonferroni’s multiple comparison test. Significant differences are relative to cells with no POS (***P<0.001, *P<0.05), or between cells treated with 50 and 200 µg/mL POS at each time point (^###^P<0.001). (**B**) AFGs were visualized by confocal immunofluorescence microscopy. Internalized rhodopsin-positive POS were detected using an anti-rhodopsin antibody (RetP1). Phalloidin staining was used to monitor cell limits and a Z-projection is represented. Arrows point to RetP1-positive POS aggregates that are attached to the apical cell membrane. Scale bar: 20 μm. (**C**) The number of intracellular AFGs were quantified using ImageJ and are represented as number of AFGs per field of view ± SEM of 3 independent experiments. (**D**) Intracellular RetP1-positive POS were quantified using ImageJ and are represented as number of RetP1-positive POS per field of view ± SEM of 3 independent experiments. Statistical comparison was performed as for panel A (***P<0.001, *P<0.05) (**E**) POS degradation was assessed using an anti-rhodopsin antibody (1D4) by Western blot analysis. Images are representative of two independent experiments.

Consistent with these results, in hfRPE AFGs were only first observed by confocal microscopy 24h after POS feeding and were still detected 3 and 7 days later (Fig. 1B), indicating that they are very stable and not easily degraded by these cells. Quantification of the number of AFGs observed per field of view, revealed a very similar pattern to that obtained using flow cytometry (Fig. 1C), showing that both techniques reliably measure the amount of AFGs in cells.

To confirm that these AFGs were indeed originating from undigested phagocytosed POS, POS internalization was monitored using an antibody that specifically recognizes the N-terminal (intradiscal) domain of rhodopsin (RetP1) ^31^, the main protein component (>90%) of the bilayer disk membranes of rod photoreceptors. As expected, immediately after POS feeding (0h), rhodopsin-positive structures were detected dispersed throughout the cytoplasm, indicating that POS were efficiently phagocytosed, while no AF was detected. Intracellular rhodopsin-positive staining decreased shortly after POS feeding (4h) and continued to decrease over time (Fig. 1B, D). Degradation of rhodopsin was also assessed by Western blot, using an antibody raised against the C-terminal (1D4 epitope) domain ^31^, since this domain is degraded in the phagocytic pathway before lysosomal fusion unlike the RetP1 epitope which is only lost after phagolysosome fusion ^31^. A band of the expected size was detected after POS feeding (0h) and was still detected 4h later (Fig. 1E) but not after 24h, a time where RetP1 labeling is still detected (Fig. 1B, C). This data suggests that early stages of phagosome processing are complete within 24h but lysosome fusion with phagosomes is still incomplete 24h post-feeding of POS.

We note that large rhodopsin-positive POS aggregates, labelled with RetP1 antibody, were found attached to the apical cell membrane 24h after POS feeding and were still detected after 7 days (Fig. 1B; Fig. S1C). We hypothesize that these rhodopsin-positive POS aggregates did not enter cells and remain strongly attached to the plasma membrane, despite extensive washes. Importantly, in most imaging experiments we also stained the RPE layer with phalloidin and analyzed only granules that were clearly within the cytoplasm.

Our results show that one single feeding of human fetal primary RPE cells with porcine POS leads to the formation of stable AFGs, mimicking the appearance of lipofuscin *in vivo*. Moreover, AFG appearance is POS-concentration dependent and increases with time. These data confirm that AF derives from POS and suggests that lysosome fusion is required for AF appearance.

### Intracellular AFGs Contain Undigested POS, are Bound by a Single Membrane and Contain Late Endosome/Lysosome markers

To confirm that AFGs result from phagolysosome formation, we performed correlative-light electron microscopy (CLEM) and studied AFGs 3 days after the POS pulse (Fig. 2A). At this time point, AFGs appear to be contained within a continuous single membrane and contain within their lumen membrane structures that resemble a more disorganized version of the regular shaped isolated segment membranes characteristic of *in retina* POS ^31^ and the isolated purified fractions of our porcine POS preparations (Fig. S1A). Very similar membrane-bound content was found in samples at 7 days after POS pulse (Fig. 2B).

**Figure 2.**
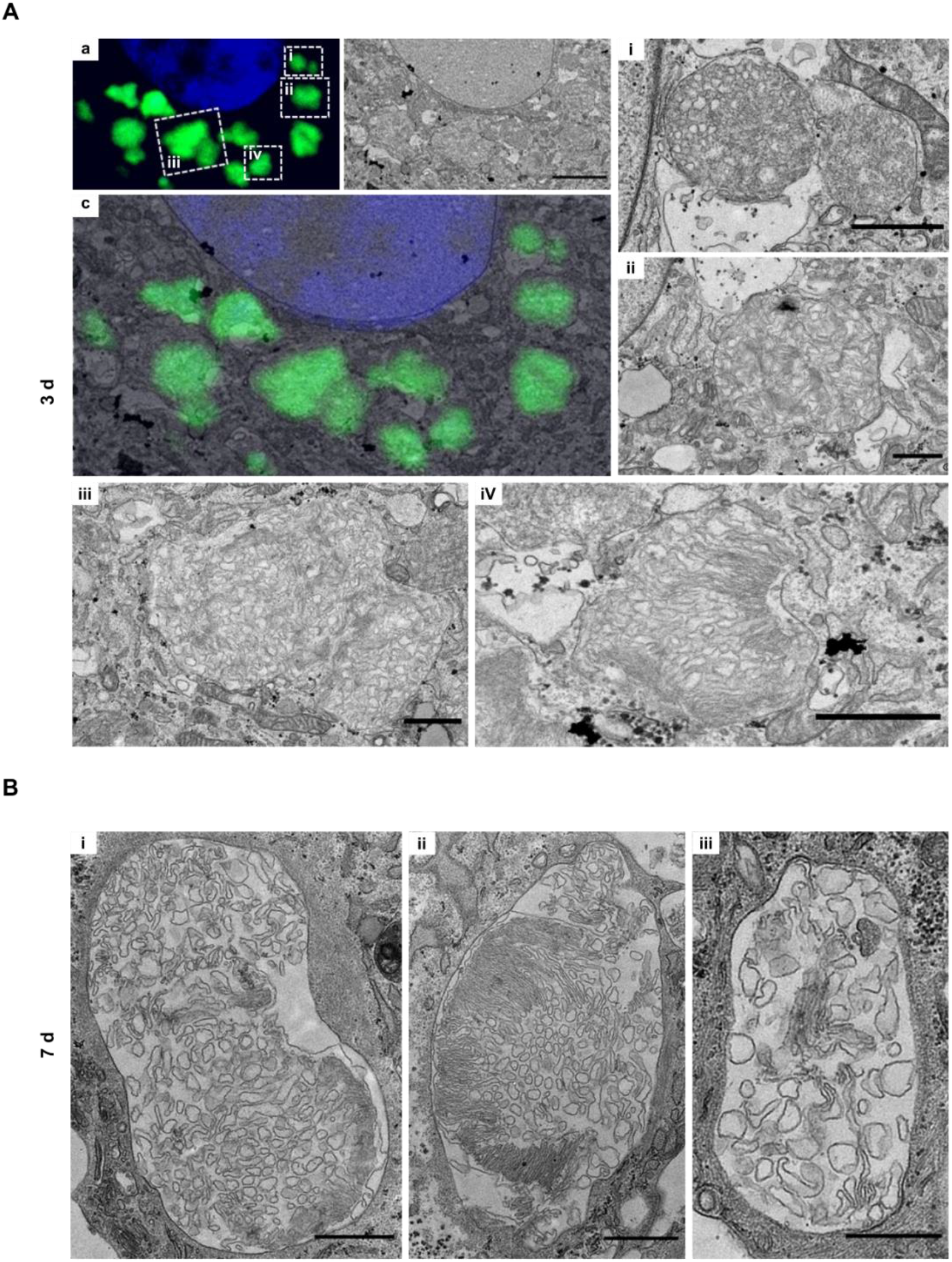
AFGs are surrounded by a single membrane and contain POS-like material. (**A**) hfRPE grown on Matek gridded dishes were fed with POS, washed and chased for 3 days. Samples were processed for CLEM as in “Materials and Methods”. a. confocal stack showing nucleus in blue and AFGs in green. b. TEM section of the equivalent area of the same cell as in a. Scale bar: 5 μm. c. overlay of a and b. i-iv. TEM insets of areas shown in a. Scale bars: 1 μm. (B) hfRPE grown on coverslips were processed for TEM 7 days after POS feeding. Examples of membrane-bound AFGs. Scale bars: i and ii, 1 μm; iii, 0.5 μm

The above results predict that AFGs contain markers of phagolysosomes ^32, 33^, Therefore, we immunostained RPE cells with the established lysosome markers, LAMP1 and CTSD (Fig. 3A, B). At 3 and 7 days post POS-feeding, approximately 70% of the intracellular AFGs were observed to be at least partially surrounded by the lysosomal membrane marker LAMP1 (Fig.3 A, C). A smaller percentage of AFGs, about 20% exhibit staining for the luminal hydrolase CTSD (Fig. 3 B, D). Interestingly, CTSD was observed to occupy the peripheral lumen of the granules surrounding the core AF.

**Figure 3.**
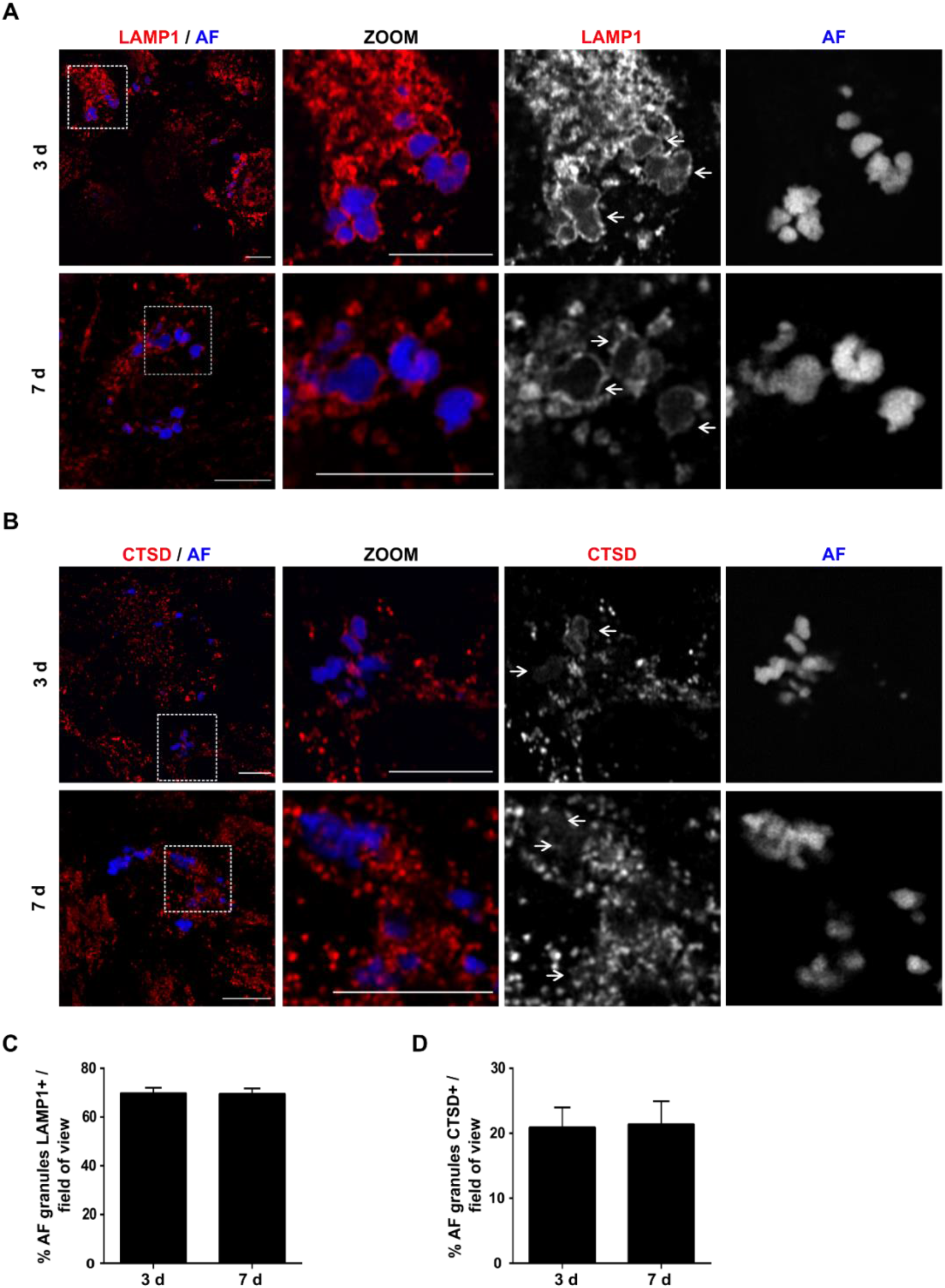
AFGs colocalize with late endosome/lysosome markers. hfRPE pulsed with 200 μg/mL POS and chased for the indicated time points were stained with (**A**) mouse anti-LAMP1 (red) or (**B**) rabbit anti-Cathepsin D antibodies (red) and AF (blue). Only a Z-slice is represented. The region outlined with a square was zoomed (ZOOM) and the different channels separated. Arrows indicate areas where AFGs are contained/surrounded by LAMP1 or Cathepsin D staining. Scale bar: 10 μm. AFGs surrounded by LAMP1 (**C**) or Cathepsin D (**D**) were quantified using ImageJ and are represented as % of granules per field of view ± SEM of 3-5 independent experiments.

We conclude that AFGs are stable over time, contain remains of ingested POS within a single membrane and contain lysosome markers, consistent with granules that evolve from phagolysosomes.

### Lysosome Fusion is Required for AFG Formation but Lysosome Dysfunction Enhances AF Accumulation

We next explored the role of lysosomes in AFG formation. We started by interfering with phagosome-lysosome fusion using a Rab7 inhibitor, CID1067700^34^. Rab7 is a Ras-like GTPase that is crucial for endo-lysosome and phago-lysosome fusion (reviewed in ^35^). Incubation with CID1067700 for 3 days significantly decreased AF levels, in a dose dependent manner, when compared with control cells in the absence of the inhibitor (Fig. 4A).

**Figure 4.**
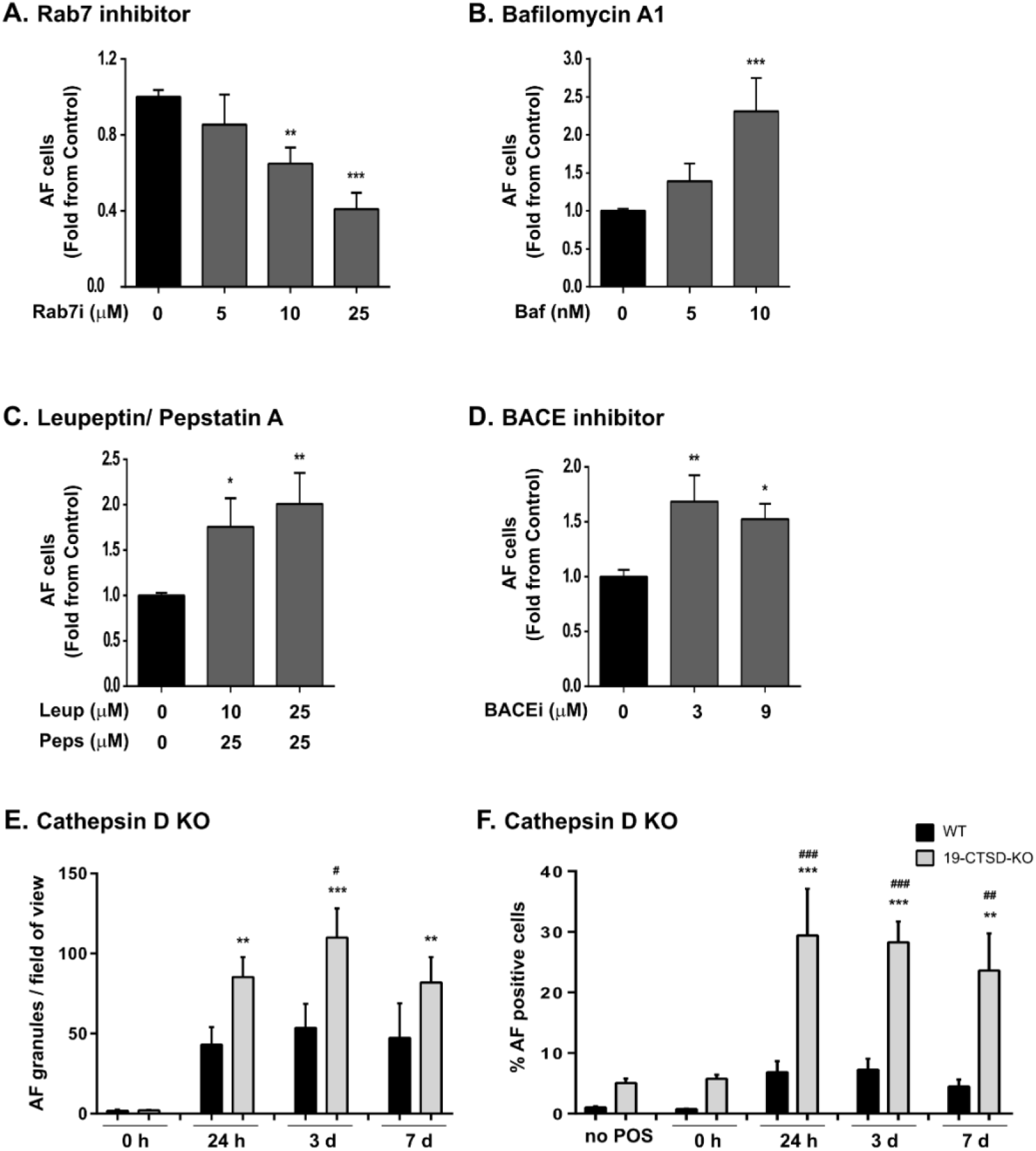
Lysosome function affects AFG formation. hfRPE were pulsed with 200 μg/mL POS and incubated for 3 days with different concentrations of (**A**) Rab7 inhibitor, (**B**) Bafilomycin A1, (**C**) Leupeptin and Pepstatin A, and (**D**) BACE inhibitor. AF levels were quantified by flow cytometry as in “Material and Methods”. Results are represented as mean ± SEM of at least 3 independent experiments. Statistical comparison was performed as in Figure 1 (***P<0.001, **P<0.01, *P<0.05). (**E**) ARPE-19 wild type (WT) and 19-CTSD-KO cells were fed with POS and chased for different time periods. Intracellular AFGs visualized by confocal immunofluorescence microscopy were quantified using ImageJ and are represented as number of AFGs per field of view ± SEM of 3 independent experiments. (**F**) ARPE-19 WT and 19-CTSD-KO AF-positive cells were quantified by flow cytometry as in “Material and Methods” at different time points after POS feeding. Results are represented as mean ± SEM of 4 independent experiments Statistical comparison was performed as in Figure 1, significant differences are relative to no POS (***P<0.001; **P<0.01) or between ARPE-19 and 19-CTSD-KO cells at each time point (^###^P<0.001, ^##^P<0.01; ^#^P<0.05).

The formation of AFGs following POS phagocytosis indicates that phagolysosome formation did not lead to complete digestion, at least in a proportion of them. The AFGs appear to contain partially digested POS by CLEM and TEM (Fig. 2), as discussed above, and the AF probably results from lipid oxidation within the phagolysosome lumen. We therefore hypothesised that the resultant AFGs are dysfunctional phagolysosomes. Therefore, we attempted to disrupt lysosomal function and observe its effects. We first fed RPE cells with POS and later treated with the compounds below to ensure that drug treatment did not interfere with POS binding or internalization.

We started by interfering with lysosomal pH using Bafilomycin A1. Bafilomycin A inhibits the lysosomal proton pump V-ATPase, thus impairs lysosome acidification and indirectly lysosomal enzyme activity ^36^. We observed that increasing concentrations of Bafilomycin A led to increasing levels of AF appearance (Fig. 4B). This suggests that, in the presence high number of of fusogenic lysosomes, an acidic lysosomal pH is critical for a complete POS digestion, and that AFGs result from incompletely digested POS.

We then interfered with hydrolytic activity of lysosomes. First, we evaluated AF formation in the presence of the protease inhibitors Leupeptin and Pepstatin A, powerful inhibitors of lysosomal enzymes. Cells incubated with increasing concentrations of these inhibitors showed increasing AF levels in RPE cells (Fig. 4C). Similar results were obtained when cells were incubated with the BACE1 inhibitor PF-7802 (Fig. 4D), described as having specific CTSD inhibitor activity as an off-target effect ^37^.

As an alternative to using drugs, which can induce undescribed off-target effects, we generated a CTSD KO ARPE-19 (19-CTSD-KO) cell line using CRISPR/Cas9 (Fig. S3A). As expected, 19-CTSD-KO no longer express the pro-form or the mature form of CTSD protein (Fig. S3B). AFGs were detected 24h after POS feeding in WT and 19-CTSD-KO cells (Fig. 4E), phenocopying the results obtained in hfRPE cells shown in Fig. 1. Quantification of the number of AFGs observed by microscopy showed that 19-CTSD-KO cells contain more AFGs in all the time points studied when compared with WT cells (Fig. 4E and S3C). The same trend was observed using flow cytometry (Fig. 4F). Interestingly, 19-CTSD-KO cells have higher basal levels of AF (at 488 nm wavelength) in the absence of POS when compared with WT cells (Fig. 4F), possibly because cargo degradation is chronically impaired in these cells, thus generating AF.

Unlike hfRPE, adult primary pRPE monolayers contain a lysosomal system that was previously operating within an intact retina and therefore presumably contain fully operational lysosomes. Nevertheless, these cells also produce AFGs after treatment with a single pulse of POS in a time-dependent manner (Fig. S2A, B). Since these are adult cells, some AFGs are present before POS feeding, presumably acquired during the lifetime of the animal from which they were derived. To distinguish between POS-derived AFGs and previously developed AFGs we loaded the POS with gold (Fig. S4) and performed CLEM so that AFGs containing POS-derived gold could be identified. Some intense AFGs appeared to lack gold particles so may have been pre-formed, although we cannot exclude the presence of gold particles in another section plane (Fig. 5A). In addition, less intense AFGs contained POS-derived gold, in agreement with the data from hfRPE, indicating that the AFGs are POS-derived. The gold particles in the AFGs are clearly aggregated, indicating that the POS are at least partly broken down, liberating the gold particles that they contain, which have aggregated in the acidic lumen of the lysosome (Fig. 5A). Our data with hfRPE indicated that, although delivery of POS to lysosomes is necessary for AF production, lysosome dysfunction and hence incomplete degradation of POS, increased AF production. We have found that the pRPE have limited tolerance to inhibitors of lysosome function. Instead, we mildly UV-irradiated POS prior to feeding to the cells in order to inhibit POS degradation. Treatment of ARPE19 cells with UV-irradiated POS has previously been shown to induce accumulation of AF and we found that although uvPOS pretreatment had no clear effect in hfRPE, it clearly elevated AF production in adult pRPE (Fig. S2D). CLEM of pRPE treated with gold loaded UV-irradiated POS showed that intense AFGs contained non-aggregated gold particles, indicating incomplete degradation of POS and/or elevation of lysosomal pH (Fig. 5B). A comparison of cells treated with a single pulse of POS or uvPOS and chased for 5 days clearly shows that uvPOS-derived gold accumulates in larger granules where it remains monodisperse, compared to the smaller granules containing aggregated POS-derived gold (Fig. 5C). Thus, using a different approach to targeting the lysosome directly, we have provided further evidence that incomplete POS degradation enhances AFG formation.

**Figure 5.**
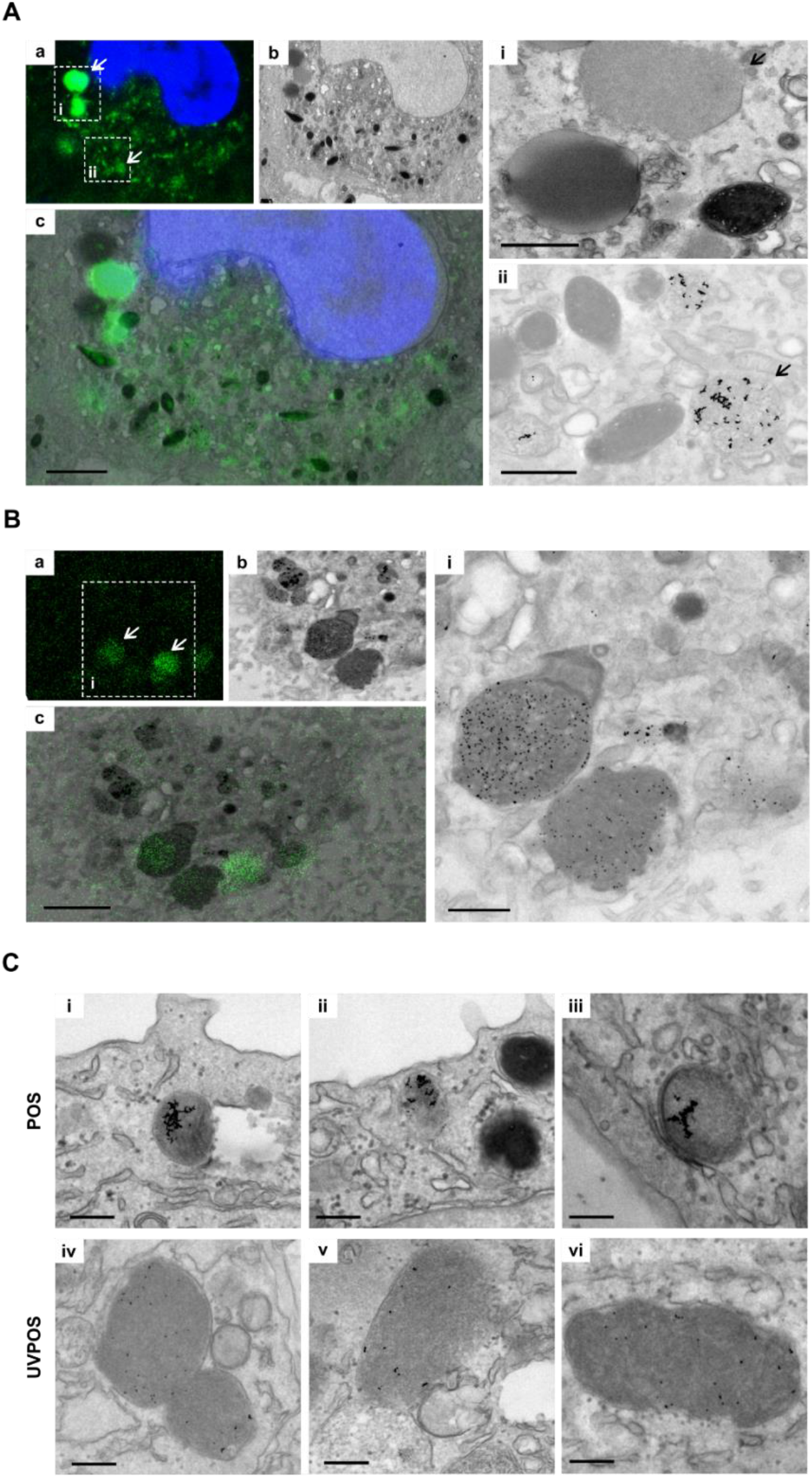
AFGs contain undigested gold-labeled POS in pRPE. pRPE grown on Matek gridded dishes were fed with (**A**) gold-tagged POS or (**B**) UV-irradiated POS (UVPOS), washed and chased for 3 days. Samples were processed for CLEM as in “Materials and Methods”. a. confocal stack showing nucleus in blue and AFGs in green. b. TEM section of the equivalent area of the same cell as in a. c. overlay of a and b. Scale bar: 2.5µm. i and ii. TEM insets of areas shown in a. Scale bars: 1 μm. (**C**) pRPE grown on coverslips were processed for TEM 5 days after feeding with gold-tagged POS or UVPOS. The presence of gold in persistent AFGs confirms that they are derived from POS. Scale bar: 500 nm.

## Discussion

The present results confirm previous studies suggesting that AFG derive from phagocytosed POS and further suggest that some lysosome activity, in particular phagosome-lysosome fusion is required for AFGs formation. However, lysosome dysfunction, particularly loss of CTSD activity, leading to incomplete digestion of POS is required for the appearance of AF and the formation of stable AFGs. Therefore, decreased lysosomal activity leads to less degradative capacity, reduced POS degradation and increased AFGs accumulation.

Previous studies have reported that repeated phagocytosis of POS leads to the progressive accumulation of AFGs within cultured primary RPE cells and RPE cell lines ^19–25^. Here we optimized a method which leads to the accumulation of AFGs after a single feeding of POS (or UV-irradiated POS), presumably due to the high concentration of the POS material. Unlike other tissues (notably neurons), RPE AFGs have a known source and their appearance can be induced by feeding RPE with POS, representing a dose and time-dependent assay. As AFGs are surrogates of RPE lipofuscin, a marker of cell stress and disease, this assay allows a unique opportunity to study the mechanisms of formation of AFGs, a process that remains ill-characterised. Here we confirm previous studies that suggest that AFGs/lipofuscin derive from POS phagocytosis ^19–25, 29, 30^. An inverse correlation was observed between rhodopsin staining and AF (Fig. 1). In fact, as rhodopsin-staining of phagosomes decreased, AF progressively developed. Consistently, gold particles phagocytosed within POS, and liberated as POS are degraded, are found within AFGs. AF is detected by 24h after POS feeding and progressively increases over 7 days (Fig. 1). At 3 and 7days post-feeding, the AFGs looked similar and exhibited partially digested POS surrounded by a single membrane (Fig. 2). These results suggest that AFGs have not lysed and released their content into the cytoplasm. Furthermore, a single membrane was always observed on the periphery of these granules at the time points studied. Lipofuscin studied in intact retinas are similarly bounded by a single membrane but contain a homogeneous lumen ^14^. This difference is presumably due to the much longer maturation time of lipofuscin compared to AFGs observed in this study. Similar single membrane-bound AFGs have been reported in RPE-J cells ^38^, suggesting that this is indeed a process of phagocytosis and not autophagy.

Having established the source of AFGs from phagosomes, we asked whether AFGs exhibited characteristics of phagolysosomes. We stained AFGs with two widely used markers of lysosomes, the lysosome membrane marker LAMP1 which clearly stained the limiting membrane of a majority of AFGs and the luminal enzyme CTSD shown to occupy the peripheral lumen of the granules surrounding the core AF in only around 20% of AFGs (Fig. 3). These results confirmed that AFGs have resulted from the fusion of lysosomes to form phagolysosomes and further suggest that CTSD is consumed within AFGs while LAMP1 is retained at the surface of the granules. A recent study followed the fate of POS and oxidatively modified POS (oxPOS) in ARPE-19 cells for 72 h and found that POS were sequentially trafficking to LAMP1 and LAMP2 as reported here but in that study the use of fluorescently-labelled POS prevented the analysis of formation of AFGs ^32, 33, 38^. In post-mortem RPE derived from AMD patients, enlarged and annular LAMP1-positive compartments were observed ^39^, which could represent AFGs even though AF was not studied.

Several reports have also shown LC3 colocalization with internalized fluorescently-labelled POS ^32, 33, 38^, as LC3-associated phagocytosis contributes to efficient POS degradation and recycling of bis-retinoids (38). However, we were unable to colocalize LC3 with AFGs either formed in hfRPE or in ARPE-19 cells (results not shown). This could be due to technical limitations related to the weak anti-LC3 signal obtained in our cells, under our testing condition. Alternatively, only a proportion of phagosomes are reported to stain for LC3 at a given time ^40^, thus AFGs may form from LC3-negative phagosomes or have lost LC3 during the maturation of AFGs.

The formation of phagolysosomes (or endolysosomes) normally leads to a complete digestion of the phagosome material, with reformation of “terminal” lysosomes ^41^. Whilst some POS-phagosomes will be fully digested, the formation of AFGs indicates that at least a proportion is not. The AFGs appear to contain partially digested POS by CLEM and TEM (Fig. 2), as discussed above, and the AF probably results from lipid oxidation within the phagolysosome lumen, suggesting that they represent dysfunctional lysosomes, as observed in many pathological conditions, from lysosome storage diseases to age-related neurodegenerative diseases ^15^. We show here that classical ways of inducing lysosomal dysfunction lead to increased AF in RPE cells. In fact, disruption of lysosomal pH (Bafilomycin A) or hydrolytic activity (CTSD, Leupeptin/Pepstatin) promote increased intensity of AF within AFGs and/or increased numbers of AFGs. Additionally, prevention of full degradation of POS by mild UV-irradiation of the POS also increased AF in adult pRPE. In agreement with the present studies, ammonium chloride, an agent that increase lysosomal pH and therefore serves as a lysosomal inhibitor, induced the appearance of AF in RPE even in the absence of POS and an 8-fold increase in AF after continuous POS feeding^42^. Conversely, acidification of lysosomes via acidic nanoparticles and pharmacological agents reduce AF in RPE cells ^20, 29, 30^.

In summary, the present study establishes that lysosomal dysfunction is a major contributor in AFG formation. These observations suggest that therapies targeted towards improvement of lysosome function, *eg*, via decrease of pH as proposed by Mitchell and colleagues ^29, 30^ or lysosome rejuvenation, *eg*, via autophagy activation are promising avenues to delay the progression of intermediate or dry AMD, currently an important medical unmet need.

## Acknowledgments

We would like to thank the Cell Culture facility at CEDOC, the scientific and technical assistance of T. Pereira, C. Andrade and A. Farinho from the CEDOC Flow Cytometry, Microscopy and Histology facilities, A.L. Sousa and E.M. Tranfield at the Instituto Gulbenkian de Ciência Electron Microscopy Facility, for technical expertise and help with EM optimisation and processing.

## SUPPLEMENTARY FIGURE LEGENDS

**Figure S1.**
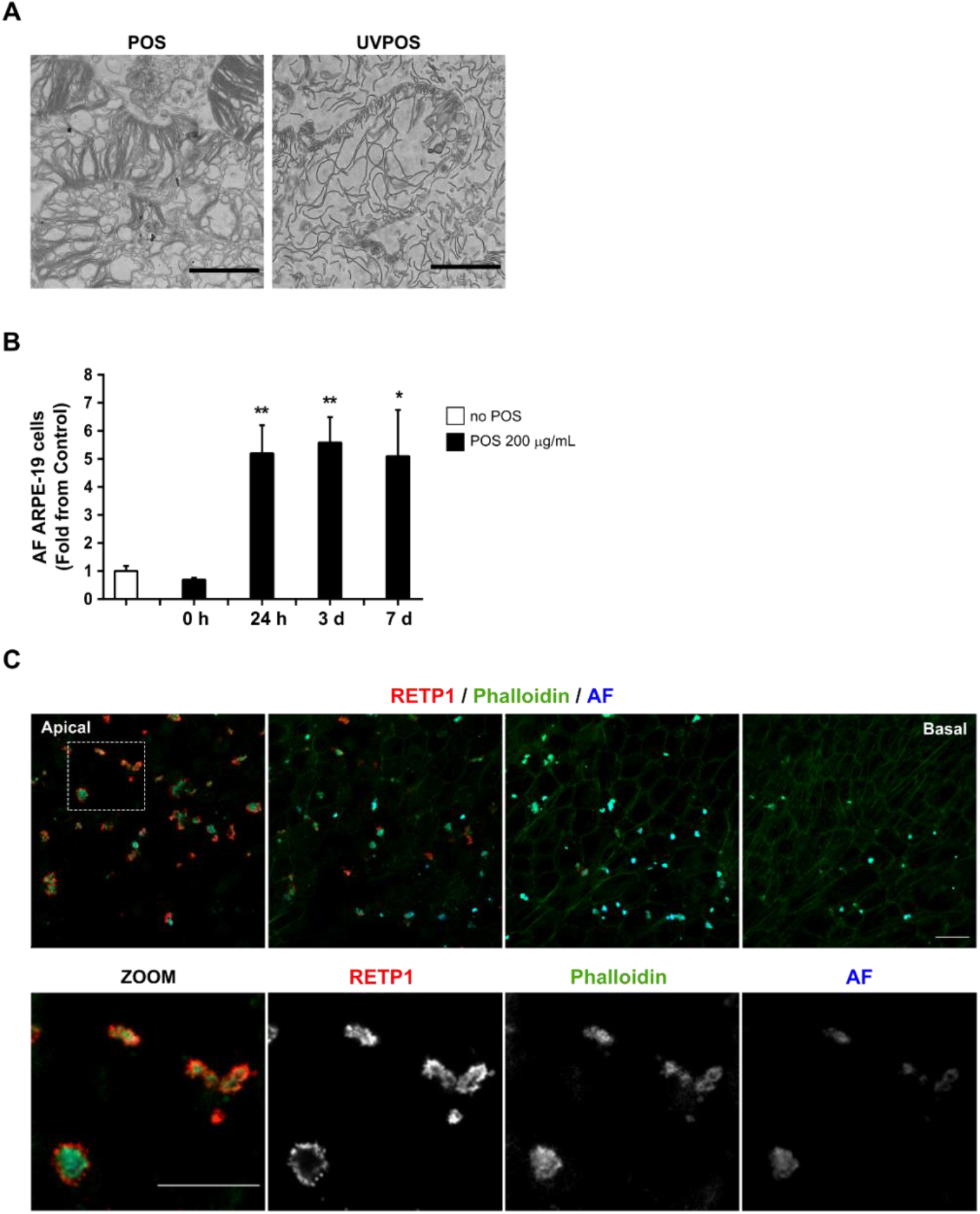
Kinetics of AF appearance in different RPE cell lines. (**A**) Porcine POS and UV-irradiated POS were isolated as described in “Material and Methods”. Pellets were processed for EM and random sections of the pellet were imaged. Scale bar: 2 μm. (**B**) ARPE-19 cells pulsed with 200 μg/mL POS were monitored at the indicated time points after POS incubation. AF levels were quantified by flow cytometry as in “Material and Methods” and are represented as mean ± SEM of 3 independent experiments. Statistical comparison was performed using one-way ANOVA followed by Dunnet’s multiple comparison test, significant differences are relative to cells with no POS (**P<0.01; *P<0.05). (**C**) hfRPE cells were pulsed with 200 μg/mL, chased for 3 days and AFGs were visualized by confocal immunofluorescence microscopy (blue). Rhodopsin-positive POS were detected using an anti-rhodopsin antibody (RetP1, red) and Phalloidin staining (green) was used to monitor cell limits. Different Z-slices are represented from the basal to apical cell layer. The region outlined with a square was zoomed (ZOOM) and the different channels separated. Scale bar: 20 μm.

**Figure S2.**
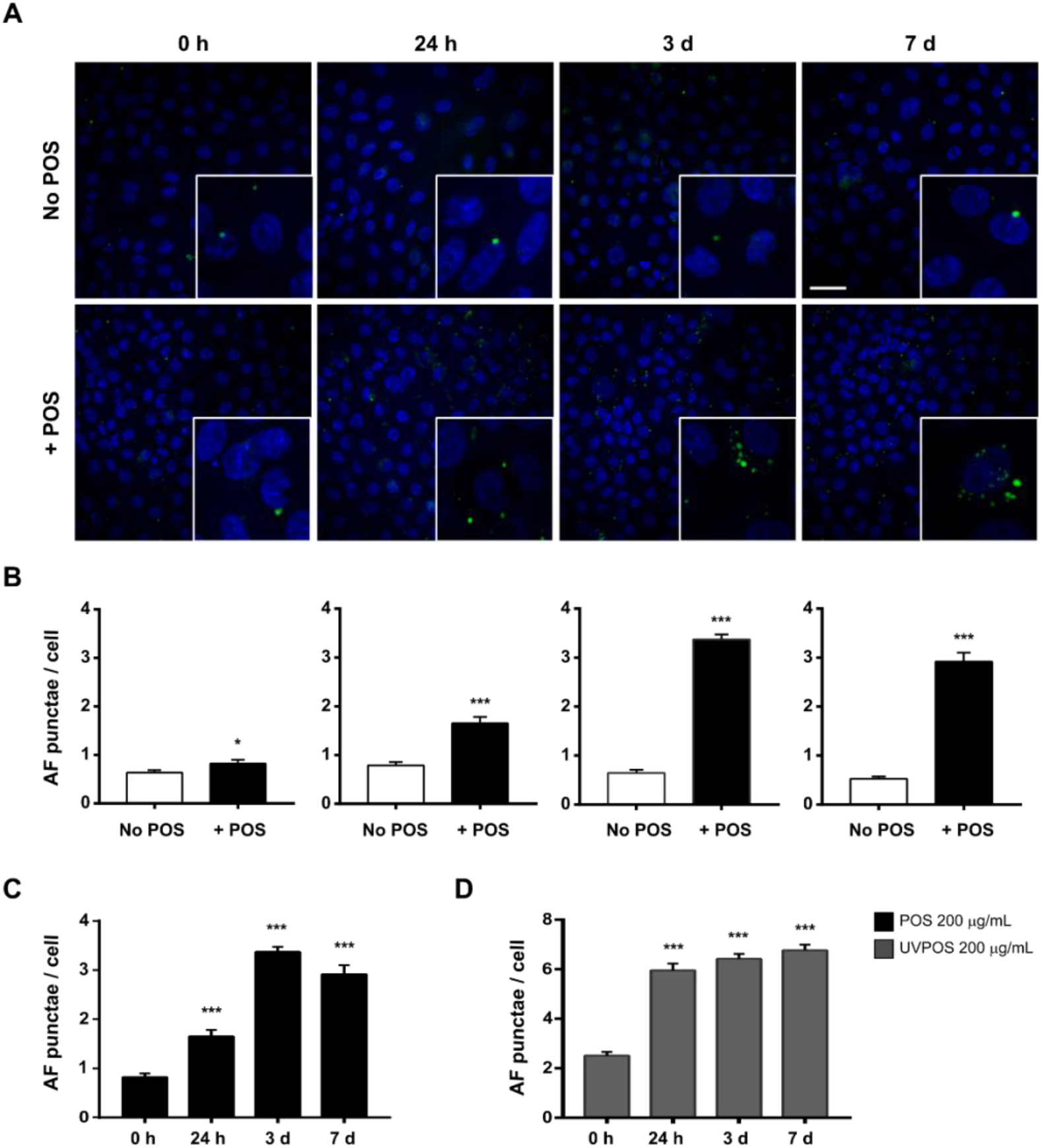
Quantification of AFGs in pRPE. pRPE were pulsed once with POS or UV-irradiated POS and chased for the indicated time points before fixing and imaging AF in the 488 emission channel with a high-content imaging platform. (**A**) Representative maximum intensity projections of confocal slices are shown. Blue: DAPI; Green: AF. Scale bar: 15 μm (**B**) The difference in the number of granules per cell between cells with no POS and POS, at the different time points, and the number of granules per cell overtime in (**C**) cells incubated with POS or (**D**) UV-irradiated POS (UVPOS) were quantitated from the z-stacks. Statistical comparison was performed using unpaired Student t test or ANOVA followed by Dunnet’s multiple comparison test; significant differences are relative to cells with no POS or POS 0h (****P<0.001; *P<0.05).

**Figure S3.**
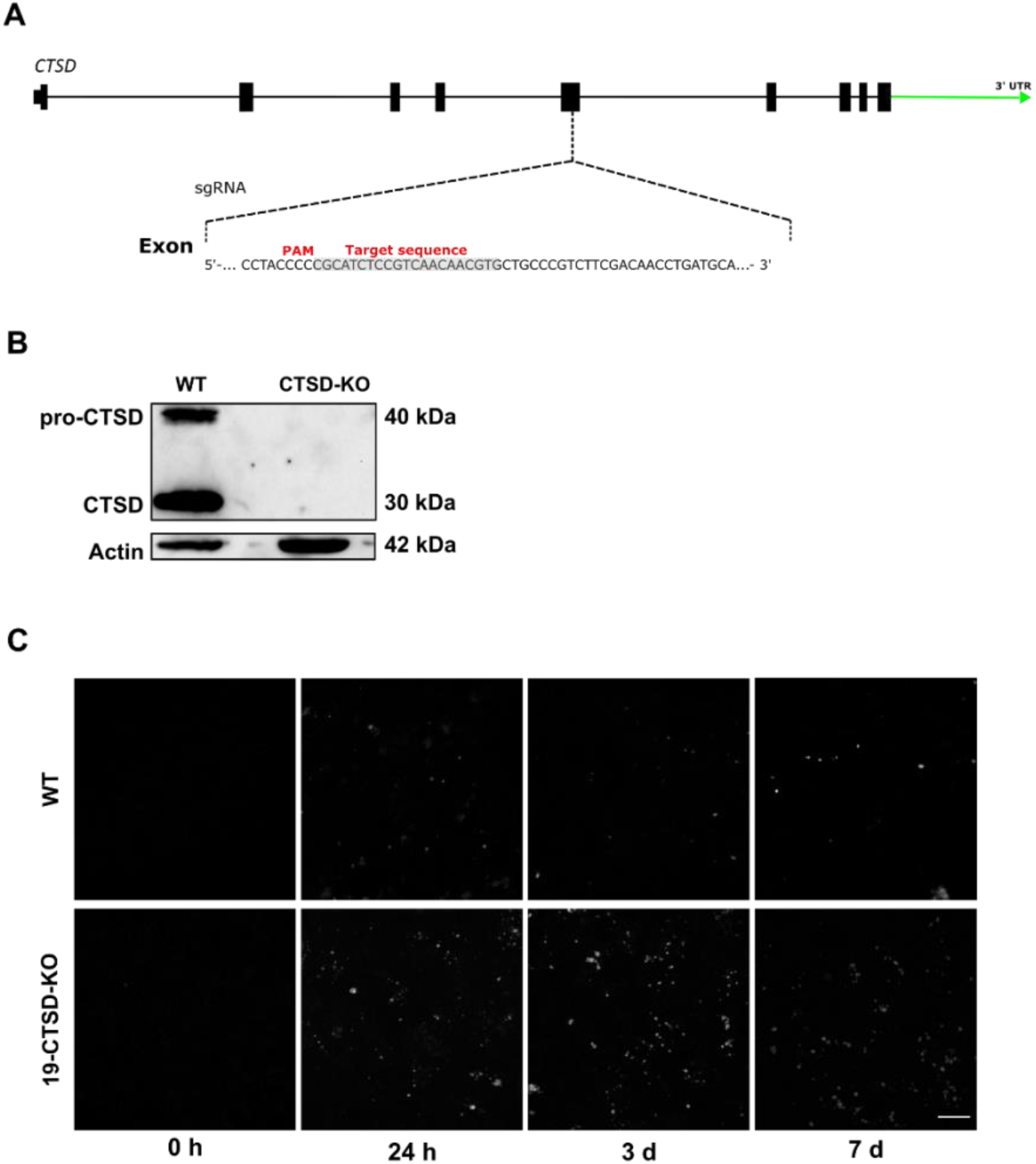
CTSD KO ARPE-19 cell line validation and AFG formation. **(A**) Schematic representation of the CTSD^-/-^ CRISPR/Cas9 strategy targeting at CTSD exon 4. The scheme represents the wild-type endogenous human chromosome sequence. The sgRNA target sequence is highlighted in light grey and the PAM sequence is in red, on the fourth exon. (**B**) CTSD protein levels in ARPE-19 wild-type (WT) and mutant (CTSD^-/-^) cells assessed by western blot. (**C**) Representative confocal immunofluorescence microscopy images of AFG in ARPE-19 wild type and 19-CTSD-KO cells feed with POS and chased for different time periods. Scale bar: 20 μm.

**Figure S4.**
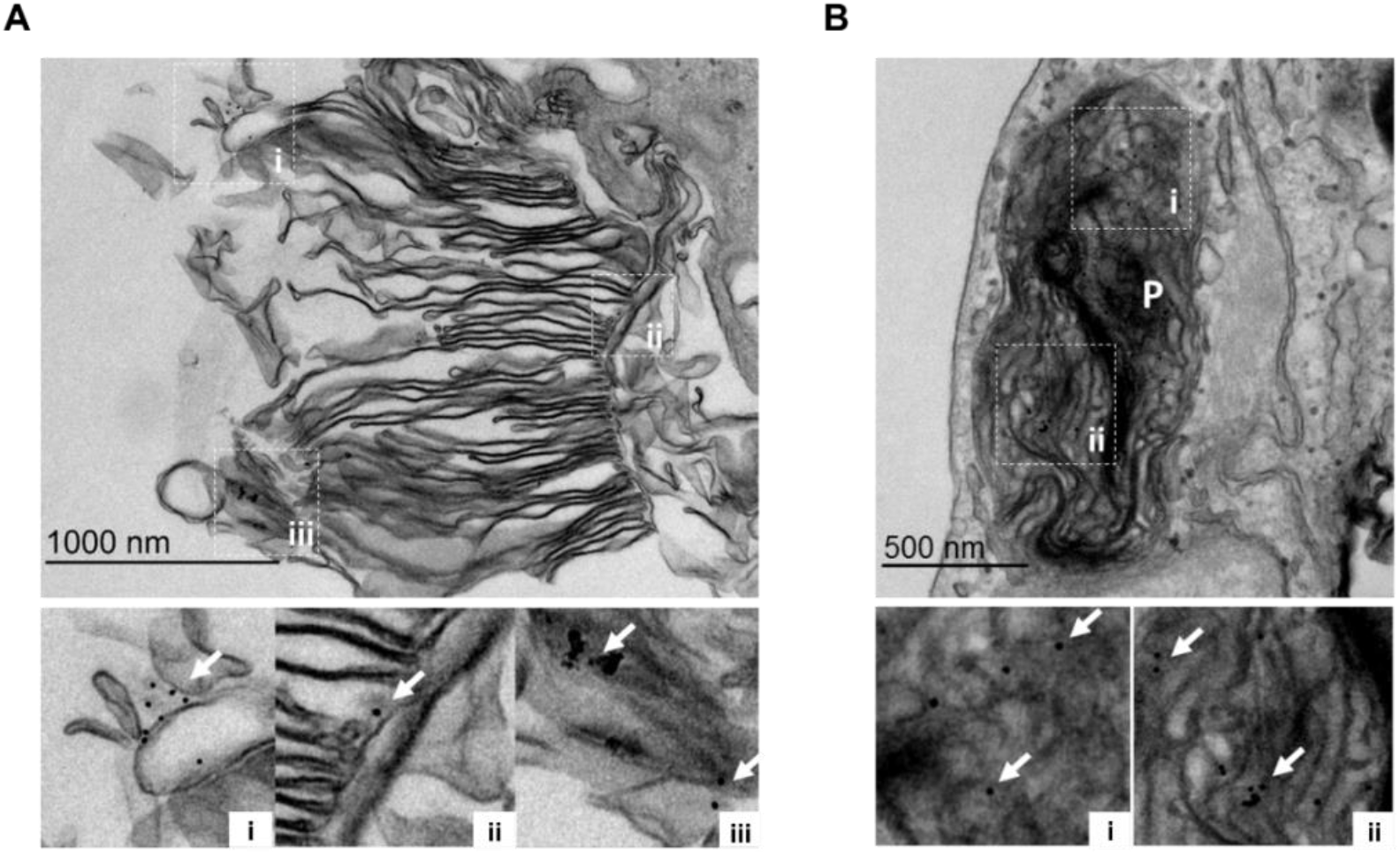
POS phagocytosis tracking by gold loaded POS with TEM. POS were tagged with 10nm gold colloid by sonication before feeding to pRPE cells in a 4-hour pulse observation by conventional EM. (**A**) At the end of the pulse, POS on the cell surface (pre-internalization) are clearly tagged with gold indicated by the arrows. (**B**) 4h post-pulse, phagosomes within the RPE are also identifiable by the presence of gold.

